# A saga of agonistic interactions in house crickets (*Acheta domesticus*): A direct and indirect effects perspective

**DOI:** 10.1101/2025.01.28.635358

**Authors:** M. A. Sekhar, Brooke A. Rothamer, Erin Gillam, Ned A. Dochtermann

## Abstract

Agonistic behaviours are widely observed across multiple taxa and are critical in shaping hierarchies, influencing resource acquisition, survival, and reproductive success. Individuals often alter their behaviour in response to the traits of others, referred to as indirect effects. Individuals also differ in their mean behavioural responses, referred to as direct effects. The combination of indirect and direct effects produce the observed social interactions. Importantly, these effects are typically measured on isolated parts of the full sequence of behaviors and traits expressed during social interactions. Here, we used house crickets, *Acheta domesticus*, to investigate how direct and indirect effects shape behaviors and traits across an agonistic interaction. We found that the probability of initiating aggression, but not contact, was influenced by both direct and indirect effects independent of mass. Amplitude, peak frequency, and pulse duration of stridulations occurring during agonistic interactions were influenced by direct effects but were not strongly influenced by indirect effects. These results demonstrate that the strength of indirect and direct effects vary over the course of an agonistic interaction and differentially affect the specific components of these interactions. Understanding when and how these effects are important is necessary for understanding agonistic behaviour and its evolution.

## Introduction

Social groups allow for cooperation in resource acquisition, reproduction, protection from predators, and more [1]. The formation and persistence of groups are driven by selective pressures that benefit the individuals involved through stable social relationships [1,2]. Within social groups, individuals will both cooperate and compete with conspecifics [3]. While cooperation is essential for group dynamics, including resource acquisition and protection from predators, competition within groups occurs because of limited resources, such as food, space, and mating opportunities [4].

One common way competition manifests among individuals is via agonistic behaviour [5]. Agonistic behaviour includes actions related to conflict, aggression, and submission. These actions can cause harm, or represent threats of harm, during social interactions [6]. This behaviour has been observed in both intra- and intersexual interactions. For example, in Asian agamid lizards (*Phrynocephalus vlangalii*), female–female aggression serves dual functions of mate and resource defence, especially during mating seasons and in the presence of familiar males [7]. While less commonly described, intersexual aggression is also frequently observed. For example, female harlequin frogs (*Atelopus varius*) chase males from their territories and try to dislodge males that have already amplexed them [8]. Most often, agonistic interactions are described between males. In male field crickets (*Gryllus integer*), aggression is displayed through physical combat and vigorous acoustic signalling i.e., stridulation [9]. Both the physical actions and acoustic signalling that occur during agonistic actions can be influenced by phenotypic characteristics like mass and sexual maturity [9]. Generally, agonistic interactions are critical in shaping hierarchies and influence survival rates and reproductive success [6]. However, there is still much to be understood about the underlying mechanisms driving these interactions [10]. A key question is how individual’s phenotypes shape the dynamics of these interactions. In particular, how phenotypic variation influences agonistic interactions requires further exploration.

In social interactions, like those during which agonistic behaviours are exhibited, the combination of phenotypic characteristics across individuals influence social outcomes [11–13]. While much research has examined how certain phenotypic characteristics impact agonistic interactions, this is often not done within the quantitative genetics framework of direct and indirect genetic effects [11–16]. This framework is necessary, as the nature of direct and indirect genetic effects can accelerate or inhibit phenotypic evolution [14]. For a focal individual, how their genotype influences a social interaction is the direct genetic effect (DGE), while the influence of their social partner’s genotype is the indirect genetic effect (IGE) [11–16]. For example, in Mediterranean field crickets (*Gryllus bimaculatus*) individuals that are highly aggressive themselves—a direct genetic effect—elicited reduced aggression in social partners as an indirect genetic effect [17].

At the phenotypic level DGEs and IGEs combine with environmental contributions to produce direct effects (DE) and indirect effects (IE) [17,18]. Direct and indirect effects have been documented for isolated components of social behaviour in many species. For example, among dung beetles (*Onthophagus taurus*), both DEs and IEs contributed significantly to the expression of maternal care behaviours [19]. Direct effects influence maternal behaviors, such as provisioning and nurturing offspring, while indirect effects shape the environment that mothers provide for their offspring, including brood mass and resource allocation, which subsequently affect offspring phenotype [19]. These DEs and IEs also have the potential to alter evolutionary outcomes [20,21]. For example, in the Mediterranean field crickets, the negative relationship between focal and opponent behaviour could lead to evolutionary stasis [17].

Despite these examples, how DEs and IEs influence behavioural expression across the course of an agonistic interaction is unclear. Instead, the focus has been on isolated components of an interaction. Understanding the magnitude of DEs and IEs over the whole course of an interaction is important because IEs affect the rate of evolutionary change by increasing or decreasing the genetic variation available for selection [13,14,22,23]. If the specific components of an interaction (e.g. Figure 1) are differentially influenced by IEs, this changes the potential for evolutionary change across the interaction. Consequently, understanding the evolution of social behaviour generally and dyadic interactions specifically requires estimating both direct and indirect contributions over the course of an entire interaction.

**Figure 1.**
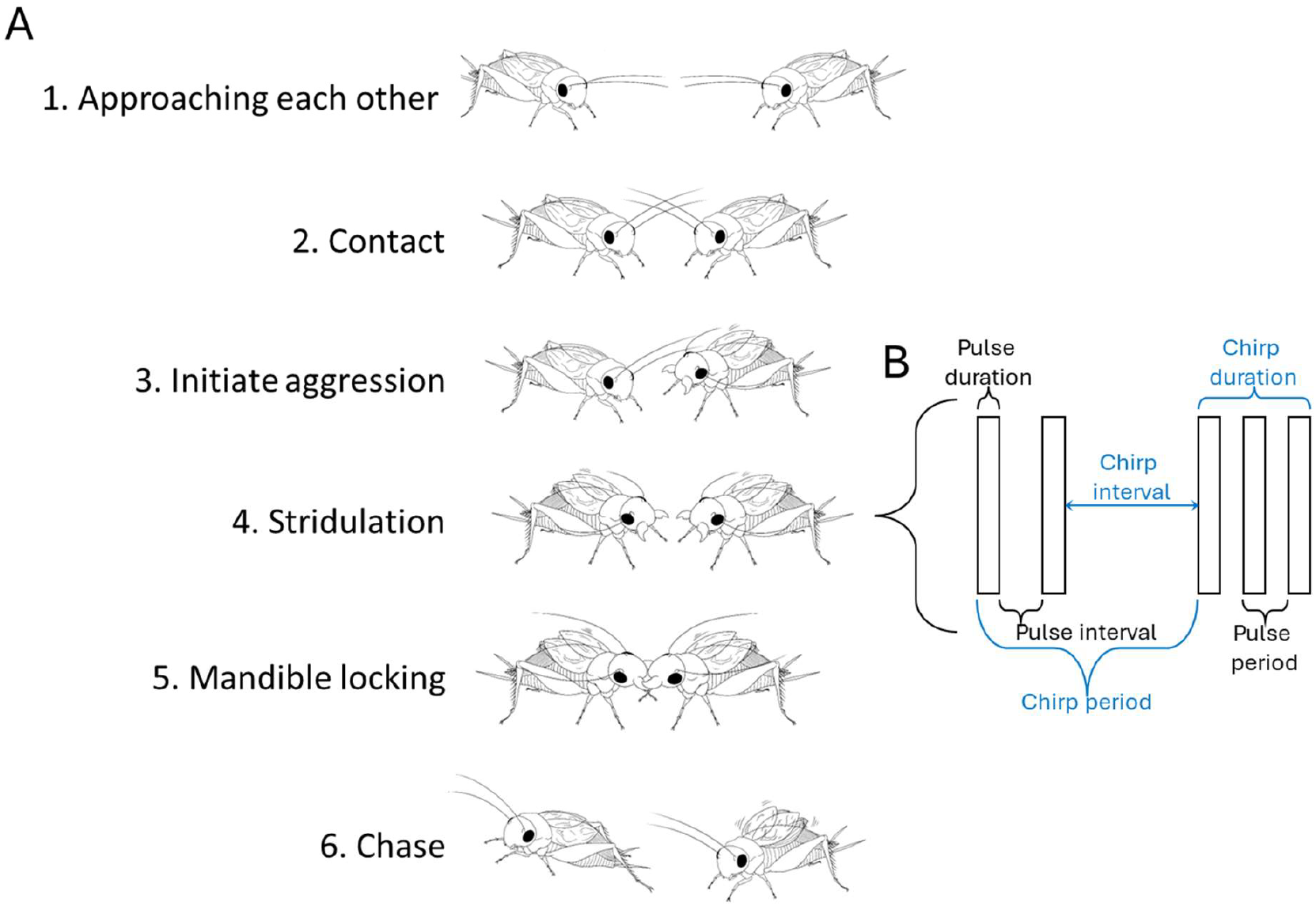
The sequence of behaviours during an agonistic interaction (A) and the parameters estimated for calls produced during this sequence (B). A) In *Acheta domesticus* an interaction begins with one or both crickets approaching each other (1) followed by contact (2), where either one or both crickets initiate antennal contact. Aggression is then initiated (3) with behaviours such as lunging, pushing, kicking, mandible locking, and aggressive stridulation. Stridulation (4), the production of aggressive calls, occurs throughout the interaction, and can also signal victory. Following this, mandible locking and repeated pushing occur (5), resulting in one cricket chasing away its opponent. After the chase, the victorious male often continues stridulating. B) Simplified oscillogram representing the temporal and spatial structure of cricket chirps. Each chirp consists of multiple pulses (represented as rectangles). We estimated spatial features amplitude and peak frequency (not shown here) and temporal characteristics chirp duration, chirp period, chirp interval, pulse duration, pulse period, pulse interval, and pulses per chirp; of each chirp. Chirp parameters are shown in blue, pulse parameters in black.

To address this lack of understanding and determine the relative contributions of direct and indirect effects on the sequence of events and associated calls during agonistic interactions, we examined aggressive contests in house crickets (*Acheta domesticus*) using a direct and indirect effects framework. *Acheta domesticus* are well known for their aggressive behaviours [24,25], which include stridulation [26]. Stridulation requires significantly less energy than other agonistic behaviours [27] but is part of an escalating sequence of responses during contests (Figure 1, [28]). Unfortunately, prior examinations of agonistic interactions in *Acheta domesticus* have primarily focused on specific components of interactions and have not estimated DEs or IEs. This prevents clear understanding of how variation among individuals contributes to dyadic interactions and their outcomes. Here we parsed agonistic interactions into constituent parts and estimated when, which part and how much of a focal individual behaviour was influenced by an opponent’s phenotype.

## Methods

To estimate direct and indirect effects on the agonistic interactions of male crickets, we conducted two experiments. The first experiment focused on the contribution of individual identities and sizes on the physical interactions between pairs. The second experiment quantified how different call parameters of aggression calls were influenced by focal and opponent identity during agonistic interactions. Jointly, these experiments allowed us to examine DEs and IEs across the course of an agonistic interaction. These experiments were conducted between June 2021 and April 2022.

### (a) Experiment 1: Estimating the strength of direct and indirect effects on the initiation of dyadic agonistic interactions

i. Cricket collection and housing: An initial *Acheta domesticus* population (n≈100) was obtained from Revier Family Farms, a vendor in Moorhead, Minnesota, USA, in May 2021. From this stock, 36 male crickets were randomly selected and housed individually in paper food containers (∼591 ml volume, ∼124 mm diameter). Crickets were provided with ad libitum food (organic chicken feed), water in small glass vials capped with cotton, and a small piece of cardboard egg carton for shelter. We maintained crickets at 27°C throughout the experiment with a 12:12 reversed dark:light cycle. Crickets were checked weekly to replace food and water and to check for maturation. Once matured, the crickets were randomly assigned into groups of four. Prior to the experimental day, crickets had their mass recorded and were randomly marked on their pronotum with one of three colours– red, yellow, green or left unmarked to distinguish individuals in interactions. The colour markings had no affect on behaviour (Table S2).
ii. Experiment 1 protocol: Behavioural trials took place between June and August 2021. Crickets were divided into nine groups of four with different colour markings within groups. For the experiment, the four crickets of a group were introduced into a rectangular arena (30cm × 15cm) separated by a partition (Figure S1). Crickets were given 5 min acclimatization time. After the acclimatization time, the partition was removed and crickets were allowed to interact freely for 15 min. All the interactions were digitally recorded using a Panasonic HC V770 and behaviours were analyzed later from the video recordings. The trials were repeated 24 hours later, a duration sufficient to eliminate carryover effects (MAS personal observation). A total of nine independent groups, with two bouts each, and four crickets per group were observed.
iii. For each trial and each of the possible pairs of individuals, we recorded contact, initiation of aggression, and chases (Figure 1A). The three behaviours were recorded and scored as follows: 1) contact – initiation of antennal contact: whichever cricket initiated contact received a score of 1 and the other received a 0. If both initiated contact at the same time, then both were scored as 1; 2) initiate aggression – initiation of aggression after contact: this included lunging, pushing, kicking, antennal locking and stridulation. Whichever cricket initiated aggression received a score of 1 and the other scored as 0. If both initiated aggression at the same time, then both were scored as 1; 3) chase – within a pair if the winner was able to successfully chase away its opponent then it was scored as 1 and opponent was scored as 0. These methods are modified from [29,30].
iv. Calculating inter- and intraobserver reliability scores

To estimate inter- and intraobserver reliability scores for each behaviour, each bout was scored from videos multiple times. For interobserver reliability, two observers (MAS and BAR) scored the videos independently, in a random order, and were blinded to group ID. For intraobserver reliability, all videos were scored twice randomly by the same observers. Reliability was estimated as the concordance correlation coefficient [31].

### (b) Experiment 2: Estimating the strength of direct and indirect effects on calling parameters

i. Cricket collection and housing: This experiment was done during two separate time periods to reach our *a priori* determined sample size of 50 males. The experiment was conducted between September to November 2021 and March to April 2022. In September 2021, a large sample of crickets were again obtained from Revier Family Farms. From these, 100 individuals were randomly selected and housed individually as in experiment 1. Once matured, 40 male crickets were randomly assigned to groups of ten males (four total groups). This was repeated in March 2022 to create a fifth group of ten males.

For identification, half of the individuals were marked with white paint on their pronotum. Individuals with the white mark were recorded as focal individuals (but see analysis description below). Unmarked individuals were recorded as opponent individuals. The colour mark had no effect on the individual or on the outcome of interactions (Table S3). Within a group, each focal individual interacted with each of the five different opponents over 10 days. Each day’s interaction represented a separate bout.

For trials, cricket pairs were moved to a square arena (20 × 20cm) and were separated by a partition (Figure S2). The floor of the arena was covered with fine playground sand to more closely represent natural conditions and to record higher quality aggressive calls (the sand floor reduced sound reflection). Crickets were given 5 min for acclimatization. After 5 min the partition was removed and the crickets interacted freely for the next 3 min. During this time, the aggressive calls produced by both focal and opponent individuals were recorded using a Tascam recorder (DR-07X, Linear PCM recorder) and videos were recorded using a Panasonic HC V770 camcorder in order to identify who called at what time. Once the trial was over crickets were returned to their original container. After 48 h, the same process was repeated but with a different opponent from the group. This gap prevented carry over effects of previous fights. Crickets do not retain memory of a novel association for long, with newly acquired memories typically decaying within 4 h [32,33]. A total of 25 interactions within a group were recorded for all groups except the fifth, in which one individual died after the third bout. We recorded a total of 123 bouts overall across all five groups. Calls were then analyzed using Raven Pro and Avisoft to estimate spatial and temporal parameters. For each call we quantified: amplitude, peak frequency, chirp duration, chirp period, chirp interval, pulse duration, pulse period, pulse interval and pulses per chirp (Figure 1B). Calls with frequencies above 7000 Hz were excluded as probable artifacts. Video recordings were processed using BORIS [34], to identify which cricket called at what time. This was then matched to estimates of call parameters.

### (c) Statistical methods

For both experiments, each interaction and each chirp were individually analyzed. In experiment 1, this resulted in analysis of 1600 total contacts, 1126 total initiations of aggression, and 26 total chases. Due to the low number of chases, this variable was excluded from subsequent analysis. In experiment 2, there were 10573 total chirps with 26277 total pulses. This led to large but varied sample sizes by individual for chirp and pulse parameters (Table S1).

For experiment 1, “focal” and “opponent” roles were assigned at random, with each individual experiencing both roles across repeated encounters to enable the partitioning of phenotypic variance in behaviour [18]. This allowed us to attribute variance to 1) the focal individual’s identity (a direct effect), 2) the opponent’s identity (an indirect effect), and 3) residual within-individual variance [18,35]. For experiment 2 sampling was based on randomly assigned focal or opponent labels but analyses were based on who called: “Caller ID” and “Opponent ID”. This allowed us to analyze the entire set of calls.

We analyzed the data using a combined trait-based approach [14] and variance components approach [36] to quantify direct and indirect effects on focal behaviour [37]. Mixed-effects models were fit with focal mass and opponent mass as fixed effects [38]. Caller ID and their social partner’s ID were included as random effects. We also included bout as a random effect in the analysis of chirps. This model structure allowed us to isolate and estimate the contribution of individual characteristics (direct effects, DE) and the contribution of characteristics of social partners (indirect effects, IE) on agonistic behaviours.

The contributions of mass to behaviour—both its direct (D mass) and indirect effects (I mass)— were estimated, as mass or size has been found to be important in prior analyses of agonistic behaviour in crickets [9]. The contributions of direct and indirect effects that are independent of mass (D var and I var) were then captured by their respective random effects. The remaining, residual variation then represents the response of focal and opponent crickets to unmeasured environmental variation [39,40].

We estimated these contributions using the R statistical language [41]. We used the rptR package [42] for calculating the proportional contribution of random effects. We then used the partR2 package [43] to estimate the contribution of specific fixed effects as the semi-partial R^2^.

Because we sought to estimate the relative contributions of direct and indirect effects across a number of the behavioural components of agonistic interactions, we fit the same statistical models to all behaviours (except bout, which could not be fit to data from experiment 1). Otherwise, models only differed based on distributional assumptions. Contact and initiation of aggression were fitted to a binomial distribution; amplitude, peak frequency, pulse duration, pulse period, pulse interval, chirp duration, chirp period and chirp interval were fitted to a normal distribution; pulses per chirp was fitted as a Poisson distribution. To allow comparison, relative contributions of direct and indirect effects are presented as variance standardized effect sizes (i.e. proportion of variance explained as unadjusted ratios, [44]) regardless of statistical “significance”. Uncertainties around these effect sizes were estimated as 95% confidence intervals via parametric bootstrapping (n = 1000) and are presented in the supplemental materials (Table S1).

## Results

In experiment 1, both intra- and inter-observer reliability scores for the behavioural trials were high. For contact and initiation of aggression, intra-observer reliability were 0.93 and 0.95, respectively. However, intra-observer reliability for chase was lower (0.70). As mentioned above, due to the few observations of this behaviour, chase was not analyzed further. Inter-observer reliability was also high for both contact (0.83) and initiation of aggression (0.88).

Initiation of aggression, amplitude, peak frequency and pulse duration exhibited substantial direct effects, with 14.8%, 19.8%, 68.6% and 31.9% of the variation, respectively, explained by focal individual ID (D var, Figure 2). Indirect effects played a substantial role in explaining variation only for initiation of aggression (18.2%). This indicates a substantial impact of opponent interactions independent of mass strictly during the initial phase of dyadic agonistic interactions.

**Figure 2.**
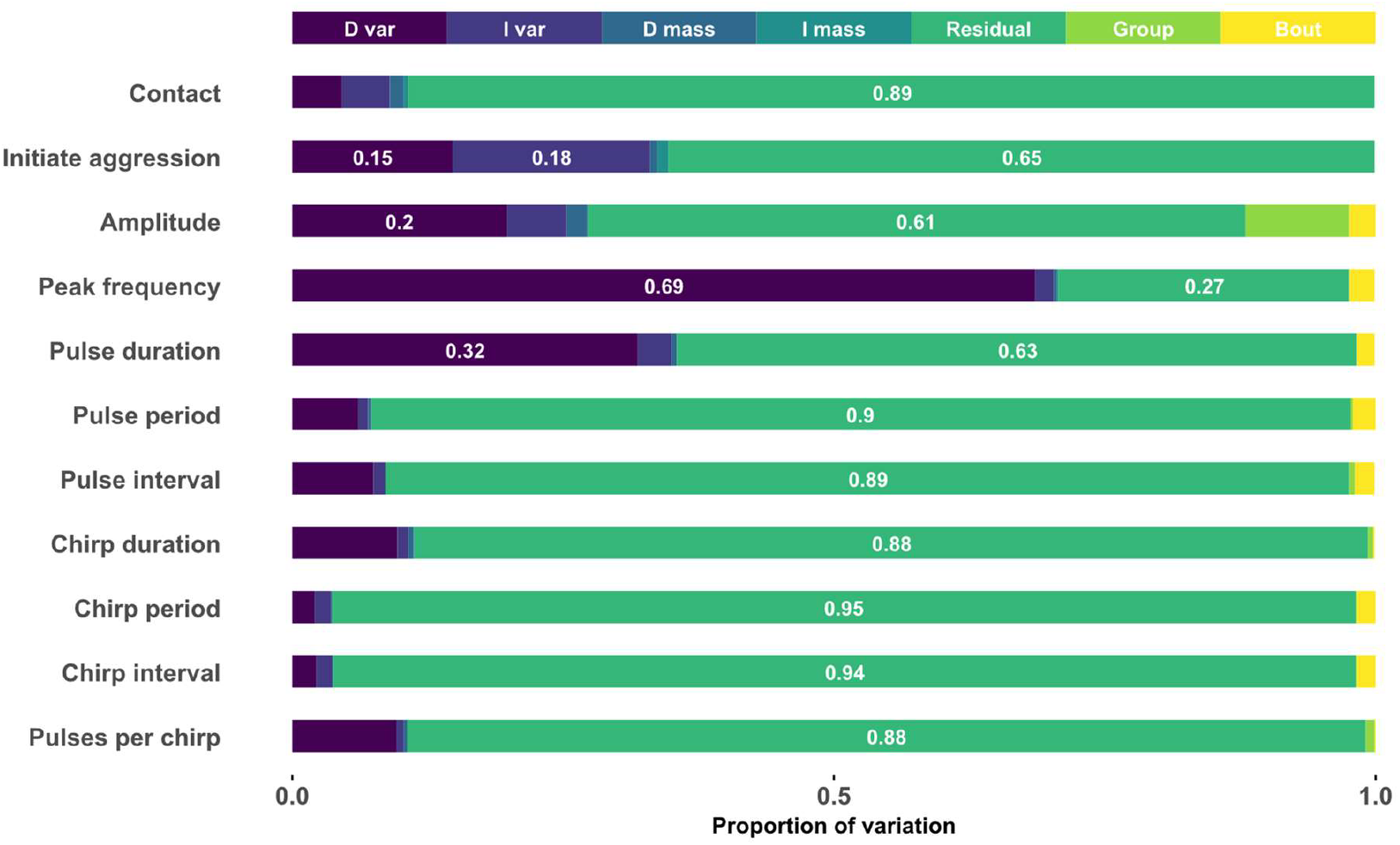
The proportion of variation explained by different factors. Each bar presents the relative contribution of: direct effects independent of mass (D var), indirect effects independent of mass (I var), direct effects due to mass (D mass), and indirect effects due mass (I mass). Also included are the contributions of residual variation, group effects, and bout effects.

Variation in most parameters was minimally influenced by mass—either as a direct or indirect effect—accounting for less than 5% of the variation among pulse duration, pulse period, pulse interval, and chirp period (Table S1).

Residual variation accounted for the largest proportion of variation in several parameters (Figure 2, Table S1). In particular, over 88% of the variation was unexplained for contact, chirp duration, chirp period, chirp interval, pulse period, pulse interval and pulses per chirp.

## Discussion

Here, we aimed to improve our understanding of agonistic interactions by determining how, when, and to what extent differences among individuals affect the behaviour of social partners across the course of a dyadic, aggressive, interaction. Specifically, we quantified the proportion of variation in behaviours explained by direct (self) and indirect (partner) effects during a dyadic agonistic interaction between house crickets.

By analyzing multiple steps of an agonistic interaction (Figure 1A), we found that contact behaviour was not strongly influenced by either a focal individual’s identity or its opponent’s identity. Biologically, it makes sense that the decision to initiate contact is not strongly influenced by individual identity: initial contact may come about during neutral exploratory behaviour rather than as an explicit aggressive response [45] and represent stochasticity in encounters. In contrast, an opponent’s influence was important in the initiation of aggression, explaining 18.2% of the variation in this behaviour (Figure 2). Put another way, whether an individual initiated aggression depended in large part on who it was interacting with. This is consistent with individuals being able to assess the competitive ability of opponents [46,47] or individuals differing in the amount of aggression they elicit in social partners [18].

We also analyzed components of the aggressive calls that occur during these agonistic interactions (Figure 1B). We found that the proportion of variation explained by direct effects varied across different components of stridulations but were generally unaffected by an opponent’s identity or mass (Figure 2, Table S1). Amplitude, peak frequency, and pulse duration exhibited particularly strong direct effects, with these components differing substantially among individuals. Such differences in these acoustic components may carry reliable information about an individual’s physical attributes or condition [48]. This suggests that these components could signal the quality or fighting ability of an individual. In contrast, for most other stridulation components, residual variation accounted for over 88% of the total variation. Consequently, unknown and unmeasured plasticity and environmental conditions primarily influence these stridulation components [40]. These could include state and resource holding potential insofar as they are variable overtime and independent of mass.

Animals use a wide variety of signals to display aggression [49–51] and these signals often play a critical role in signalling an individual’s abilities during conflicts. For example, signals can convey resource-holding potential [52] and physical strength [28]. In crickets, visual, tactile, and acoustic signals are used during agonistic interactions (Figure 1, [53]). Specific components of signals have also been found to be important during agonistic interactions and in other social interactions. For example, in Blanchard’s cricket frog (*Acris crepitans blanchardi*), males respond more aggressively to lower frequency calls, suggesting that call frequency plays a role in escalating aggression [54]. Relatedly, in the robust conehead (*Neoconocephalus robustus*), females exhibit a preference for specific pulse durations in male calls, which may indicate male quality [55].

Beyond the information that may be carried by signals, mass and physical size of an individual is often an important factor in determining the outcome of agonistic interactions both in crickets and other taxa e.g. [9,56,57]. However, we found that mass had minimal impact on agonistic interactions in *Acheta domesticus*. This was true for both individuals in a dyadic interaction. While surprising, this is consistent with some other findings: In *Acheta domesticus*, mass had little effect on fight outcomes, as its influence was often minimized by resource characteristics and differences in hunger levels [58]. Similarly, in Mediterranean crickets (*Gryllus bimaculatus*), disabling an individual’s mandibles, vision, or pairing differently sized individuals did not affect fighting behaviour and fight duration [59]. This suggests mass or physical size plays a minimal role in agonistic interactions [59].

Several studies have explored the influence of mass on stridulation parameters in *Acheta domesticus* itself. Chirps with a greater number of pulses per chirp were shown to convey male size and influence female mate choice among house crickets [48]. Similarly, courtship songs of house crickets did not directly correlate with body size or other male attributes which influence female choice [60]. House crickets aggressive songs have also been shown to be associated with body size and resource-holding potential, but they do not convey information about a male’s motivation to engage in or persist in a fight [52]. Unlike these studies, our findings indicate that mass plays a minimal roles in agonistic interactions without external motivators, such as mate choice or resource acquisition. This highlights a unique context where physical traits do not clearly influence agonistic interactions.

Rather than mass and size, are other factors generally more influential in agonistic interactions? The greater contribution of direct effects other than mass to behavioural variation during agonistic interactions, as found here (Figure 2), suggests that is the case in *Acheta domesitcus*. This seems to also be the case in other species: In European fallow deer (*Dama dama*), differences in body weight or antler length did not affect fight duration or intensity suggesting that contest outcome are influenced more by self-assessment and experience than by physical asymmetry [61].

Importantly, the influence of social partners on behavioural responses is not limited to just the dyadic interactions we measured here. For example, among fruit flies (*Drosophila melanogaster*), dyadic interactions are affected by the behaviour of other individuals in the group, a second-order indirect effect [62] and described generally as an audience effect [63]. This general phenomenon can escalate to other members within a group, resulting in higher-order effects where the actions of multiple individuals affect interactions across a group. Unfortunately, empirical data for higher-order effects in a direct-indirect effects framework is limited. As one example, sleep in group living animals, often considered as an individual behaviour is significantly influenced by social interactions [64], with conspecifics shaping sleep patterns within a group [64]. Directly applying the direct and indirect effects framework can enhance our understanding of these and other higher-level group interactions.

Using a direct-indirect effects framework [13,14,18,65] reveals important aspects of the biology of agonistic interactions. Here, we found that indirect effects of opponents only had substantial influence during the initiation of aggression. Within these agonistic interactions, we also found strong direct effects on specific stridulation components—including amplitude, peak frequency, and pulse duration. The differential expression of direct and indirect across the course of an agonistic interaction has important implications of our understanding of social behaviours.Because IGEs can enhance the evolutionary potential of traits [22], our findings emphasize the importance of separating both direct and indirect effects across the course of social interactions. The differential expression we identified means that different components of an agonistic interaction can evolve at different rates. Often, disentangling direct and indirect effects in ecological studies is challenging because they can appear at unpredictable times and after extended periods [66]. However, regardless of this difficulty, estimation of both direct and indirect contributions to social interactions is necessary to properly understand the evolution of social behaviours [35,67].

## Acknowledgements

The authors thank Hieu Le for helping in cricket rearing and husbandry. The authors thank Katy Takumi for the cricket sketches. We also thank the Environmental and Conservation Sciences program and Department of Biological Sciences at NDSU for support.

## Data Availability

All data and associated analysis code is available at: https://osf.io/t6psd/?view_only=fbcb17b6deb84871922cb373602eb3bc

## Notes

https://osf.io/t6psd/

### Competing Interest Statement

The authors have declared no competing interest.

