## Supplemental Tables and Figures for "A saga of agonistic interactions in house crickets (*Acheta domesticus*): A direct and indirect effects perspective"

**Table S1**. Estimated 95% confidence intervals for the relative contributions of direct effects independent of mass (D var), indirect effects independent of mass (I var), direct effects due to mass (D mass), indirect effects due to mass (I mass), residual variation, group effects, and bout effects across behavioral components of agonistic interactions. These estimates were obtained using parametric bootstrapping (n = 1000) and standardized effect sizes for comparison across behaviors.

| Parameter | component | R2 | l_ci | u_ci |
|---|---|---|---|---|
| **Contact (N=1600)** | D var | 0.0462274 | 0.011217 | 0.077505 |
| | I var | 0.0442932 | 0.01006 | 0.074066 |
| | D mass | 0.0133781 | 0.000344 | 0.048114 |
| | I mass | 0.0041656 | 0 | 0.039195 |
| | Residual | 0.8919357 | 0.838801 | 0.945161 |
| | Group | 8.52E-11 | 0 | 0.026029 |
| **Initiate aggression (N=1126)** | D var | 0.1483743 | 0.056771 | 0.215034 |
| | I var | 0.1822985 | 0.074824 | 0.26481 |
| | D mass | 0.0065293 | 0 | 0.077612 |
| | I mass | 0.0102986 | 0 | 0.081392 |
| | Residual | 0.6524992 | 0.544315 | 0.76888 |
| | Group | 7.86E-10 | 0 | 0.093734 |
| **Amplitude (N=26277)** | D var | 0.1979891 | 0.123851 | 0.278088 |
| | I var | 0.05513 | 0.030535 | 0.0861 |
| | D mass | 0.0187151 | 0.002161 | 0.060975 |
| | I mass | 0.0005538 | 0 | 0.04199 |
| | Residual | 0.6068557 | 0.487961 | 0.701916 |
| | Group | 0.0964653 | 0 | 0.254581 |
| | Bout | 0.0242911 | 0.01056 | 0.04217 |
| **Peak frequency (N=26277)** | D var | 0.6863938 | 0.552221 | 0.751534 |
| | I var | 0.0166728 | 0.008516 | 0.027779 |
| | D mass | 0.0034723 | 0 | 0.047459 |
| | I mass | 0.0011216 | 0 | 0.045311 |
| | Residual | 0.2691889 | 0.208057 | 0.35424 |
| | Group | 0 | 0 | 0.055732 |
| | Bout | 0.0231506 | 0.009945 | 0.041265 |
| **Pulse duration (N=26277)** | D var | 0.3193914 | 0.215144 | 0.404859 |
| | I var | 0.0310311 | 0.0177 | 0.048763 |
| | D mass | 0.0053549 | 0.000233 | 0.041401 |
| | I mass | 8.55E-05 | 0 | 0.036739 |
| | Residual | 0.6280757 | 0.540447 | 0.710119 |
| | Group | 0 | 0 | 0.040322 |
| | Bout | 0.0160614 | 0.007477 | 0.027783 |
| | D var | 0.0606408 | 0.0328 | 0.089749 |

| | | | | |
|---|---|---|---|---|
| **Pulse period (N=15702)** | I var | 0.0087128 | 0.002955 | 0.016371 |
| | D mass | 0.0025531 | 0 | 0.013857 |
| | I mass | 0.0002305 | 0 | 0.011542 |
| | Residual | 0.9044795 | 0.867238 | 0.931952 |
| | Group | 0.0020964 | 0 | 0.027611 |
| | Bout | 0.0212869 | 0.008643 | 0.036568 |
| **Pulse interval (N=15702)** | D var | 0.0750717 | 0.043435 | 0.112268 |
| | I var | 0.0109685 | 0.004267 | 0.018773 |
| | D mass | 0.0004354 | 0 | 0.008764 |
| | I mass | 0.0002239 | 0 | 0.008551 |
| | Residual | 0.8902022 | 0.84764 | 0.924424 |
| | Group | 0.0046165 | 0 | 0.032508 |
| | Bout | 0.0184818 | 0.007091 | 0.033326 |
| **Chirp duration (N=10573)** | D var | 0.0974075 | 0.058125 | 0.141378 |
| | I var | 0.0101948 | 0.003736 | 0.017999 |
| | D mass | 0.0050283 | 0 | 0.025049 |
| | I mass | 0.0006721 | 0 | 0.020869 |
| | Residual | 0.8803595 | 0.82842 | 0.918563 |
| | Group | 0.0049861 | 0 | 0.032408 |
| | Bout | 0.0013517 | 0 | 0.004475 |
| **Chirp period (N=9929)** | D var | 0.0210133 | 0.00903 | 0.036966 |
| | I var | 0.0148309 | 0.005852 | 0.026697 |
| | D mass | 4.06E-05 | 0 | 0.006665 |
| | I mass | 0.0006303 | 3.91E-05 | 0.007254 |
| | Residual | 0.9454774 | 0.921079 | 0.962399 |
| | Group | 0.0013109 | 0 | 0.01773 |
| | Bout | 0.0166966 | 0.005751 | 0.029292 |
| **Chirp interval (N=9928)** | D var | 0.0222348 | 0.008714 | 0.036499 |
| | I var | 0.0152911 | 0.005697 | 0.028245 |
| | D mass | 6.52E-05 | 0 | 0.006221 |
| | I mass | 0.0006537 | 1.67E-05 | 0.006806 |
| | Residual | 0.9437809 | 0.918417 | 0.961795 |
| | Group | 0.0013658 | 0 | 0.017918 |
| | Bout | 0.0166085 | 0.005263 | 0.029653 |
| **Pulses per chirp (N=10574)** | D var | 0.0962831 | 0.052087 | 0.132128 |
| | I var | 0.0066467 | 0.000641 | 0.012212 |
| | D mass | 0.0028332 | 0 | 0.019525 |
| | I mass | 0.0004655 | 0 | 0.017258 |
| | Residual | 0.8849609 | 0.846398 | 0.932053 |
| | Group | 0.0078196 | 0 | 0.031185 |
| | Bout | 0.0009911 | 0 | 0.004391 |

**Table S2**. Summary of fixed effects for physical interactions colour impact - Contact and Initiate Aggression, in house crickets. Estimates, standard errors, z-values, and p-values are reported for each predictor, including focal and opponent mass, and colour markings. Crickets were randomly assigned into groups of four after reaching maturity, with individual mass recorded and pronotum markings applied (red, yellow, green, or unmarked). These colour markings had no significant affect on behavior.

| **Effect** | **Category** | **Estimate** | **Std. Error** | **z value** | **Pr(>\|z\|)** |
|---|---|---|---|---|---|
| Intercept | | 0.7185 | 1.21045 | 0.594 | 0.5528 |
| Focal_Mass | | 4.06539 | 3.10386 | 1.31 | 0.1903 |
| Opp_Mass | | -3.21935 | 3.30162 | -0.975 | 0.3295 |
| Focal_ColourPink | | -0.51204 | 0.27291 | -1.876 | 0.0606 |
| Focal_ColourWhite | | -0.10583 | 0.29044 | -0.364 | 0.7156 |
| Focal_ColourYellow | | -0.17969 | 0.28502 | -0.63 | 0.5284 |
| Opp_ColourPink | | 0.17101 | 0.29685 | 0.576 | 0.5646 |
| Opp_ColourWhite | Contact | -0.21374 | 0.30368 | -0.704 | 0.4815 |
| Opp_ColourYellow | | -0.03816 | 0.30395 | -0.126 | 0.9001 |
| as.factor(Bout)2 | | -0.06469 | 0.11415 | -0.567 | 0.5709 |
| Intercept | | -0.74539 | 2.34874 | -0.317 | 0.751 |
| Focal_Mass | | 7.25282 | 6.35173 | 1.142 | 0.254 |
| Opp_Mass | | -4.69719 | 6.17862 | -0.76 | 0.447 |
| Focal_ColourPink | | 0.01342 | 0.54571 | 0.025 | 0.98 |
| Focal_ColourWhite | | -0.41825 | 0.56452 | -0.741 | 0.459 |
| Focal_ColourYellow | Initiate Aggression | 0.52764 | 0.56341 | 0.937 | 0.349 |
| Opp_ColourPink | | 0.44261 | 0.53478 | 0.828 | 0.408 |
| Opp_ColourWhite | | -0.19163 | 0.55975 | -0.342 | 0.732 |
| Opp_ColourYellow | | -0.17371 | 0.55414 | -0.313 | 0.754 |
| as.factor(Bout)2 | | 0.12085 | 0.15193 | 0.795 | 0.426 |

Signif. codes:  0 ‘***’ 0.001 ‘**’ 0.01 ‘*’ 0.05 ‘.’ 0.1 ‘ ’ 1

**Table S3**. Summary of fixed effects for call parameters across multiple metrics, including Amplitude, Peak Frequency, Pulse Duration, Pulse Period, Pulse Interval, Chirp Duration, Chirp Period, Chirp Interval and Pulses per chirp. Call parameters were modeled with respect to focal mass and opponent mass, as well as the presence of a white colour mark for identification. The presence of a white mark had no effect on individual behavior or interaction outcomes.

<table>
<tr><th>Call Parameter</th><th>Effect</th><th>Estimate</th><th>Std. Error</th><th>df</th><th>t value</th><th>Pr(>|t|)</th></tr>
<tr><td rowspan="4">Amplitude</td><td>(Intercept)</td><td>-24.8517</td><td>3.6499</td><td>71.4534</td><td>-6.809</td><td>2.56e-09 ***</td></tr>
<tr><td>Call_mass</td><td>21.8543</td><td>9.8668</td><td>44.1582</td><td>2.215</td><td>0.032 *</td></tr>
<tr><td>Opp_mass</td><td>2.4842</td><td>5.4258</td><td>35.9337</td><td>0.458</td><td>0.650</td></tr>
<tr><td>Call_ID_colourWhite</td><td>-1.0396</td><td>0.7163</td><td>59.2705</td><td>-1.451</td><td>0.152</td></tr>
<tr><td rowspan="4">Peak Frequency</td><td>(Intercept)</td><td>4202.31</td><td>376.42</td><td>82.78</td><td>11.164</td><td><2e-16 ***</td></tr>
<tr><td>Call_mass</td><td>815.75</td><td>1149.11</td><td>79.23</td><td>0.71</td><td>0.480</td></tr>
<tr><td>Opp_mass</td><td>262.17</td><td>223.56</td><td>35.12</td><td>1.173</td><td>0.249</td></tr>
<tr><td>Call_ID_colourWhite</td><td>18.06</td><td>87.01</td><td>43.54</td><td>0.208</td><td>0.836</td></tr>
<tr><td rowspan="4">Pulse Duration</td><td>(Intercept)</td><td>0.0062557</td><td>0.0043904</td><td>62.0703</td><td>1.425</td><td>0.159</td></tr>
<tr><td>Call_mass</td><td>0.0131995</td><td>0.0130041</td><td>51.832</td><td>1.015</td><td>0.315</td></tr>
<tr><td>Opp_mass</td><td>0.000813</td><td>0.0044523</td><td>34.1393</td><td>0.183</td><td>0.856</td></tr>
<tr><td>Call_ID_colourWhite</td><td>-0.0001644</td><td>0.0009003</td><td>51.9244</td><td>-0.183</td><td>0.856</td></tr>
<tr><td rowspan="4">Pulse Period</td><td>(Intercept)</td><td>0.05004</td><td>0.007691</td><td>53.51</td><td>6.506</td><td>2.7e-08 ***</td></tr>
<tr><td>Call_mass</td><td>0.0325</td><td>0.02204</td><td>39.37</td><td>1.474</td><td>0.148</td></tr>
<tr><td>Opp_mass</td><td>0.00502</td><td>0.009679</td><td>25.4</td><td>0.519</td><td>0.608</td></tr>
<tr><td>Call_ID_colourWhite</td><td>0.0004132</td><td>0.001471</td><td>45.19</td><td>0.281</td><td>0.780</td></tr>
<tr><td rowspan="4">Pulse Interval</td><td>(Intercept)</td><td>0.04539</td><td>0.008732</td><td>56.21</td><td>5.198</td><td>2.91e-06 ***</td></tr>
<tr><td>Call_mass</td><td>0.01296</td><td>0.02504</td><td>43.02</td><td>0.517</td><td>0.607</td></tr>
<tr><td>Opp_mass</td><td>0.006288</td><td>0.01084</td><td>27.84</td><td>0.58</td><td>0.567</td></tr>
<tr><td>Call_ID_colourWhite</td><td>-4.477e-06</td><td>0.001678</td><td>49.24</td><td>-0.003</td><td>0.998</td></tr>
<tr><td rowspan="4">Chirp Duration</td><td>(Intercept)</td><td>0.012946</td><td>0.048584</td><td>54.41</td><td>0.266</td><td>0.791</td></tr>
<tr><td>Call_mass</td><td>0.212371</td><td>0.141531</td><td>43.12</td><td>1.501</td><td>0.141</td></tr>
<tr><td>Opp_mass</td><td>0.056164</td><td>0.05562</td><td>27.48</td><td>1.01</td><td>0.321</td></tr>
<tr><td>Call_ID_colourWhite</td><td>-0.007245</td><td>0.009366</td><td>46.51</td><td>-0.774</td><td>0.443</td></tr>
<tr><td rowspan="4">Chirp Period</td><td>(Intercept)</td><td>2.7398</td><td>1.3671</td><td>45.07</td><td>2.004</td><td>0.0511 .</td></tr>
<tr><td>Call_mass</td><td>-0.4159</td><td>3.3989</td><td>24.13</td><td>-0.122</td><td>0.9036</td></tr>
<tr><td>Opp_mass</td><td>-2.8446</td><td>2.6928</td><td>31.51</td><td>-1.056</td><td>0.2988</td></tr>
<tr><td>Call_ID_colourWhite</td><td>0.1495</td><td>0.2544</td><td>37.14</td><td>0.588</td><td>0.5602</td></tr>
<tr><td rowspan="4">Chirp Interval</td><td>(Intercept)</td><td>2.7174</td><td>1.3911</td><td>45.63</td><td>1.953</td><td>0.0569 .</td></tr>
<tr><td>Call_mass</td><td>-0.6148</td><td>3.4692</td><td>24.48</td><td>-0.177</td><td>0.8608</td></tr>
<tr><td>Opp_mass</td><td>-2.8938</td><td>2.7249</td><td>31.67</td><td>-1.062</td><td>0.2963</td></tr>
<tr><td>Call_ID_colourWhite</td><td>0.1576</td><td>0.2593</td><td>37.71</td><td>0.608</td><td>0.5469</td></tr>
<tr><td></td><td>(Intercept)</td><td>0.45647</td><td>0.33136</td><td></td><td>1.378</td><td>0.168</td></tr>
</table>

<table>
<tr><td rowspan="4">Pulses per chirp</td><td></td><td></td><td></td><td></td><td>(Z-value)</td><td>Pr(>|z|)</td></tr>
<tr><td>Call_mass</td><td>0.99752</td><td>0.97014</td><td></td><td>1.028<br>(Z-value)</td><td>0.304<br>Pr(>|z|)</td></tr>
<tr><td>Opp_mass</td><td>0.33325</td><td>0.34257</td><td></td><td>0.973<br>(Z-value)</td><td>0.331<br>Pr(>|z|)</td></tr>
<tr><td>Call_ID_colourWhite</td><td>-0.06939</td><td>0.06296</td><td></td><td>-1.102<br>(Z-value)</td><td>0.270<br>Pr(>|z|)</td></tr>
</table>

Signif. codes: 0 ‘***’ 0.001 ‘**’ 0.01 ‘*’ 0.05 ‘.’ 0.1 ‘ ’ 1

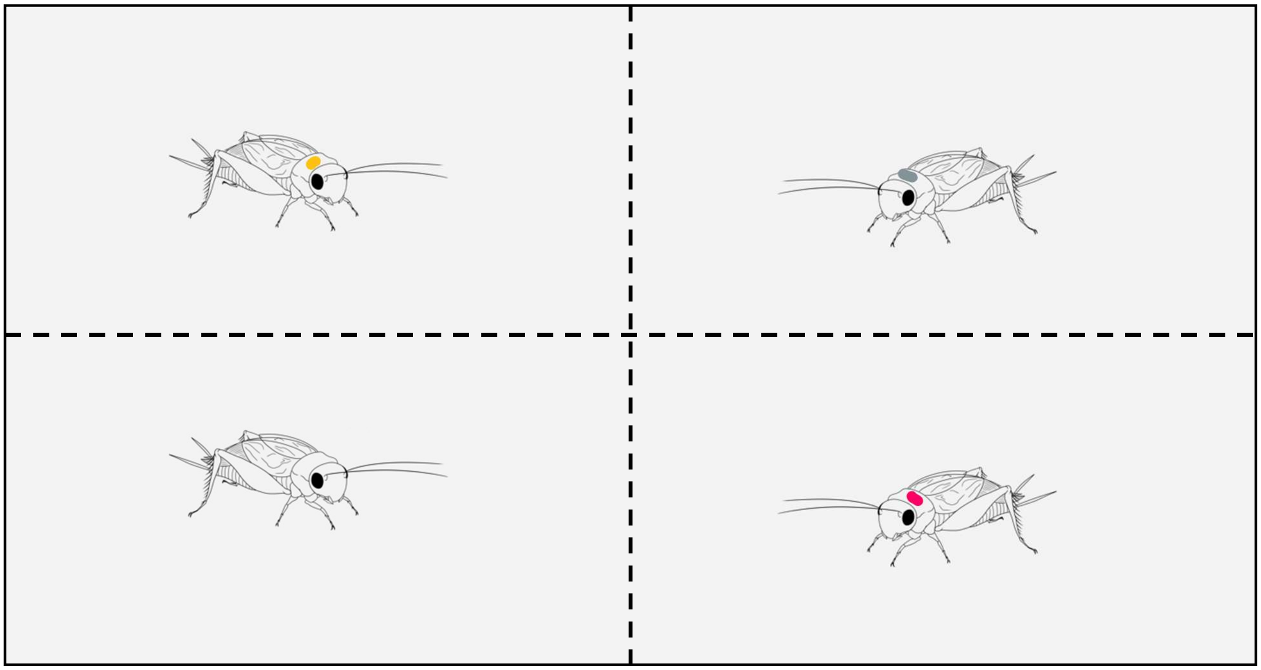

**Figure S1.** Experimental setup for behavioral trials in a rectangular arena (30 cm × 15 cm), showing four crickets (not to scale) within separate compartments, each marked with a unique colour (yellow, white, pink, and unmarked) for individual identification. Cricket were given 5 min. acclimatization time. After that, the partitions was lifted and crickets were allowed to interact freely for 15 min.

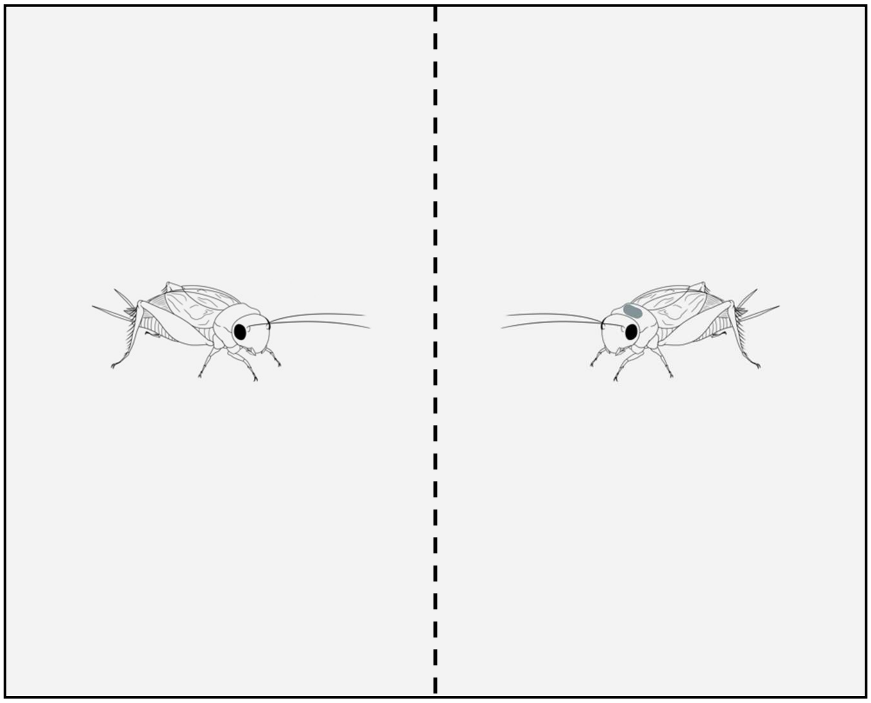

**Figure S2.** Experimental setup for dyadic trials in a square arena (20 cm × 20 cm), featuring a soil floor to mimic natural conditions and reducing sound reflection for higher-quality aggressive call recordings. Cricket pairs (not to scale) were initially separated by a partition and given 5 minutes for acclimatization. After the acclimatization period, the partition was lifted, allowing the crickets to interact freely for 3 minutes.
